# Probing differences in gene essentiality between the human and animal adapted lineages of the *Mycobacterium tuberculosis* complex using TnSeq

**DOI:** 10.1101/2021.09.21.461195

**Authors:** Amanda J Gibson, Ian J Passmore, Valwynne Faulkner, Dong Xia, Irene Nobeli, Jennifer Stiens, Sam Willcocks, Dirk Werling, Bernardo Villarreal-Ramos, Brendan W Wren, Sharon L Kendall

**Author notes:** present address (Amanda J Gibson, Valwynne Faulkner, Bernardo Villarreal-Ramos).

## Abstract

Members of the Mycobacterium tuberculosis complex (MTBC) show distinct host adaptations, preferences and phenotypes despite being >99% identical at the nucleic acid level. Previous studies have explored gene expression changes between the members, however few studies have probed differences in gene essentiality. To better understand the functional impacts of the nucleic acid differences between *Mycobacterium bovis* and *Mycobacterium tuberculosis* we used the Mycomar T7 phagemid delivery system to generate whole genome transposon libraries in laboratory strains of both species and compared the essentiality status of genes during growth under identical *in vitro* conditions. Libraries contained insertions in 54% of possible TA sites in *M. bovis* and 40% of those present in *M. tuberculosis*, achieving similar saturation levels to those previously reported for the MTBC. The distributions of essentiality across the functional categories were similar in both species. 527 genes were found to be essential in *M. bovis* whereas 477 genes were essential in *M. tuberculosis* and 370 essential genes were common in both species. CRISPRi was successfully utilised in both species to determine the impacts of silencing genes including *wag31*, a gene involved in peptidoglycan synthesis and *Rv2182c*/*Mb2204c*, a gene involved in glycerophospholipid metabolism. We observed species specific differences in the response to gene silencing, with the inhibition of expression of *Mb2204c* in *M. bovis* showing significantly less growth impact than silencing its ortholog (*Rv2182c*) in *M. tuberculosis*. Given that glycerophospholipid metabolism is a validated pathway for antimicrobials, our observations suggest that target vulnerability in the animal adapted lineages cannot be assumed to be the same as the human counterpart. This is of relevance for zoonotic tuberculosis as it implies that the development of antimicrobials targeting the human adapted lineage might not necessarily be effective against the animal adapted lineage. The generation of a transposon library and the first reported utilisation of CRISPRi in *M. bovis* will enable the use of these tools to further probe the genetic basis of survival under disease relevant conditions.

## Introduction

*Mycobacterium bovis* and *Mycobacterium tuberculosis* are closely related members of the Mycobacterium tuberculosis complex (MTBC). Although both species are >99% identical at the nucleotide level each species shows distinct host tropisms. *M. bovis*, the animal adapted species, is the main causative agent of bovine tuberculosis in cattle (1) while *M. tuberculosis* is the main cause of human tuberculosis (TB) and is responsible for ∼1.5 million deaths annually (1,2). *M. bovis* exhibits a broader host range than *M. tuberculosis* and is also able to cause TB in humans through zoonotic transfer, representing a serious public health risk in countries without a control programme in domestic livestock (2,3). The WHO recognises that zoonotic transfer of tuberculosis threatens the delivery of the end TB strategy, highlighting the importance of understanding the differences between the two species (3).

Many studies have explored the genotypic and phenotypic differences between *M. tuberculosis* and *M. bovis* in order to better understand host preference. Genome sequencing of the reference strains (H37Rv and AF2122/97) showed that the main genetic differences between these pathogens were several large-scale deletions, or regions of difference (RD), and over 2,000 single-nucleotide polymorphisms (SNPs) (4– 7). More recently, studies that include clinically circulating strains have confirmed that all animal adapted lineages share deletions RD7, 8, 9, and 10 (8). Transcriptomic studies which have measured significant changes in gene expression between H37Rv and AF2122/97 have provided a functional insight into the impacts of some of these polymorphisms (9–11). For instance, a SNP in *rskA* (*Mb0452c*) an anti-sigma factor in *M. bovis*, prevents repression of *sigK* activity, leading to constitutively high levels of expression of *mpb70* and *mpb83*, genes that encode key immunogenic antigens; MPB70 and MPB83 (12,13). Recent studies have shown that MPB70 mediates multi-nucleated giant cell formation in *M. bovis* infected bovine macrophages, but not in *M. bovis* (or *M. tuberculosis*) infected human macrophages, providing insight into bacterial effectors of the species-specific response (14). Transcriptomic studies have also indicated a differential response to *in vitro* mimics of host stresses such as acid shock and highlight the impact of SNPs in the signalling and response regulons in two-component systems such as PhoPR and DosSRT (15–18).

Genome-wide transposon mutagenesis coupled with next-generation sequencing (TnSeq) has allowed genome wide predictions of gene essentiality in *M. tuberculosis* (19– 24). These studies have provided information on the genetic requirements for *in vitro* growth under a number of conditions and also for growth in disease relevant models such as macrophages (20). Most of these studies performed in the MTBC have used strain H37Rv. More recently Tnseq of different clinical strains of *M. tuberculosis* has shown that there are strain specific differences in fitness associated with Tn insertions in certain genes. The implication of this observation is that different strains can show different antibiotic sensitivities as a result (25). To date, there has been a single reported Tnseq study performed in *M. bovis* (AF2122/97) which focused on intra-cellular genetic requirements (26)

A direct comparison of gene essentiality in *M. bovis* and *M. tuberculosis* has not been reported. Therefore, we created dense transposon libraries in both *M. bovis* (AF2122/97) and *M. tuberculosis* (H37Rv) generated on the same medium to enable direct comparisons between the two related species. We identified that there are key differences in essentiality in *M. bovis* compared to *M. tuberculosis*. We used CRISPRi to directly demonstrate that silencing the expression of a gene annotated to be involved in glycerophospholipid metabolism has different impacts on growth in the two species. This has implications for target discovery programmes as it implies that inhibition of therapeutically relevant pathways may have different impacts in the different species. This is important in the context of zoonotic tuberculosis.

## Materials and Methods

### Bacterial Strains and Culture Methods

*M. bovis* AF2122/97 was maintained on modified Middlebrook 7H11 solid medium containing 0.5% lysed defibrinated sheep blood, 10% heat inactivated foetal bovine serum and 10% oleic acid-albumin-dextrose-catalase (OADC) (27). Liquid cultures of *M. bovis* were grown in Middlebrook 7H9 medium containing 75 mM sodium pyruvate, 0.05% Tween^®^80 and 10% albumin-dextrose-catalase (ADC). *M. tuberculosis* H37Rv and *Mycobacterium smegmatis* mc^2^155 were maintained on Middlebrook 7H11 solid medium supplemented with 0.5% glycerol and 10% OADC. Liquid cultures were grown in Middlebrook 7H9 medium supplemented with 0.2% glycerol, 0.05% Tween^®^80 and 10% ADC unless stated otherwise. MycomarT7 Phagemid was propagated on *M. smegmatis* mc^2^155 lawns grown on Middlebrook 7H10 solid medium supplemented with 0.5% glycerol and 10% OADC in a 0.6% agar overlay. The strains and plasmids used or made in this study are given in Table 1.

**Table 1.**
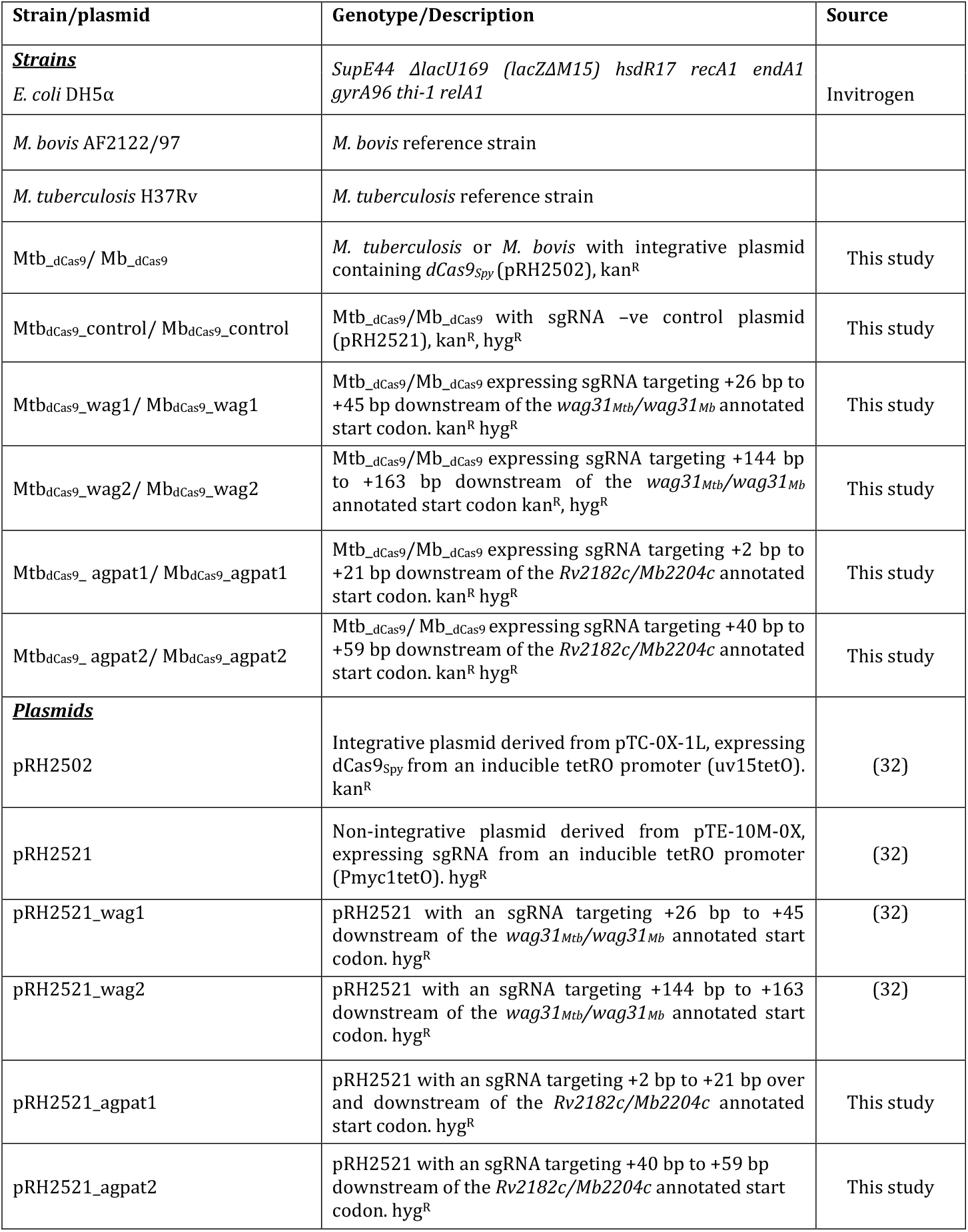
Strains and plasmids used in this study.

### Generation of Transposon Libraries

Transposon libraries in *M. bovis* (AF2122/97) and *M. tuberculosis* (H37Rv) were generated using the previously described MycomarT7 phagemid system as per Majumdar *et al* with modifications (28). Briefly, 50 ml cultures of *M. bovis* and *M. tuberculosis* at OD_600_≅ 1 were washed twice with MP buffer (50 mM Tris-HCl, pH 7.5, 150 mM NaCl, 10 mM MgSO_4_ and 2 mM CaCl_2_) at 37 °C, and then incubated with ∼ 10^11^ pfu of φMycoMarT7 phage for 16-18 h at 37 °C without rolling. Transduced bacteria were washed in pre-warmed PBS + 0.05% Tween^®^80 to remove extra-cellular phage and plated on Middlebrook 7H11 solid medium containing 0.5% lysed defibrinated sheep blood, 10% heat inactivated foetal bovine serum, 10% OADC, 25 μg/ml kanamycin and 0.05% Tween^®^80. Cultures were allowed to grow for 5-6 weeks. Concurrent CFU plating was performed to estimate transduction efficiency. Approximately 15-20 colonies from each library were used for validation of random insertion using a nested PCR strategy followed by Sanger sequencing, method and data are shown in Supplementary File S1. Libraries were scraped from the plates and incubated in liquid medium at 37°C with hourly vortexing for 3 h to homogenise. Homogenised mutants were distributed to cryovials and stored at -80°C for further selection or gDNA extraction.

### DNA Extraction

Unless stated otherwise, reagents were acquired from Sigma Aldrich. Genomic DNA from harvested libraries was isolated by a bead beating procedure (mechanical lysis) or using de-lipidation followed by enzymatic lysis as previously described by Long *et al* 2015 (29) and Belisle *et al* 2009 (30). Briefly, for mechanical lysis, library aliquots were disrupted using 0.1 mm glass beads and bead-beating by 3 × 15 sec bursts (5000 rpm) interspersed with 2 min on ice using a beat-beater (Biospec). For enzymatic lysis, libraries were de-lipidated with equal volumes chloroform-methanol (2:1) for 1 h with agitation every 15 min, suspension was centrifuged at 3,488 x g for 10 min the bacterial pellet allowed to dry for 2 h after removal of both solvent layers. De-lipidated bacteria were suspended in TE buffer and incubated with 100 μg/ml lysozyme in the presence of 100 mM TrisBase (pH 9.0) at 37°C for 12-16 h. Bacterial lysis was completed by incubating for 3 h at 55°C in the presence of 1% SDS and 100 μg/ml proteinase K (NEB). Lysates from both methods were extracted twice with equal volumes of phenol– chloroform–isoamyl alcohol (25:24:1). The aqueous layer was harvested by centrifugation at 12,000 x g for 30 min and DNA was precipitated with 0.1 volumes of 3M sodium acetate (pH 5.2) and one volume of ice-cold isopropanol overnight at -20°C. DNA pellets were washed several times in ethanol. DNA was re-suspended in water and quantity and quality were determined using a DeNovix Spectrophotometer (DeNovix Inc, USA), agarose gel electrophoresis and fluorometry using Qubit4 (Invitrogen).

### Library Preparation for Transposon Directed Inserted Sequencing

2 μg of extracted DNA libraries were resuspended in purified water and sheared to approximately 550 bp fragments using a S220 focussed-ultrasonicator (Covaris), according to the manufacturer’s protocol. Sheared DNA was repaired using NEBNext blunt-end repair kit (New England Biolabs) and purified using Monarch PCR clean-up kit (New England Biolabs). Blunted DNA was A-tailed using NEBNext dA-tailing kit and column-purified. Custom transposon sequencing adaptors, or “TraDIS tags”, (Table 2) were generated by heating an equimolar mix of adaptor standard primer and adaptor P7+index to 95°C for 7 min and then allowed to cool to room temperature. Adaptors were ligated to A-tailed library fragments using NEBNext quick ligase kit. Transposon-containing fragments were enriched by PCR using ComP7 primers ComP5 using Phusion DNA polymerase (New England Biolabs) in a 20-cycle reaction. Library fragments were subsequently cleaned up with AMPureXP purification beads (Beckman).

**Table 2.**
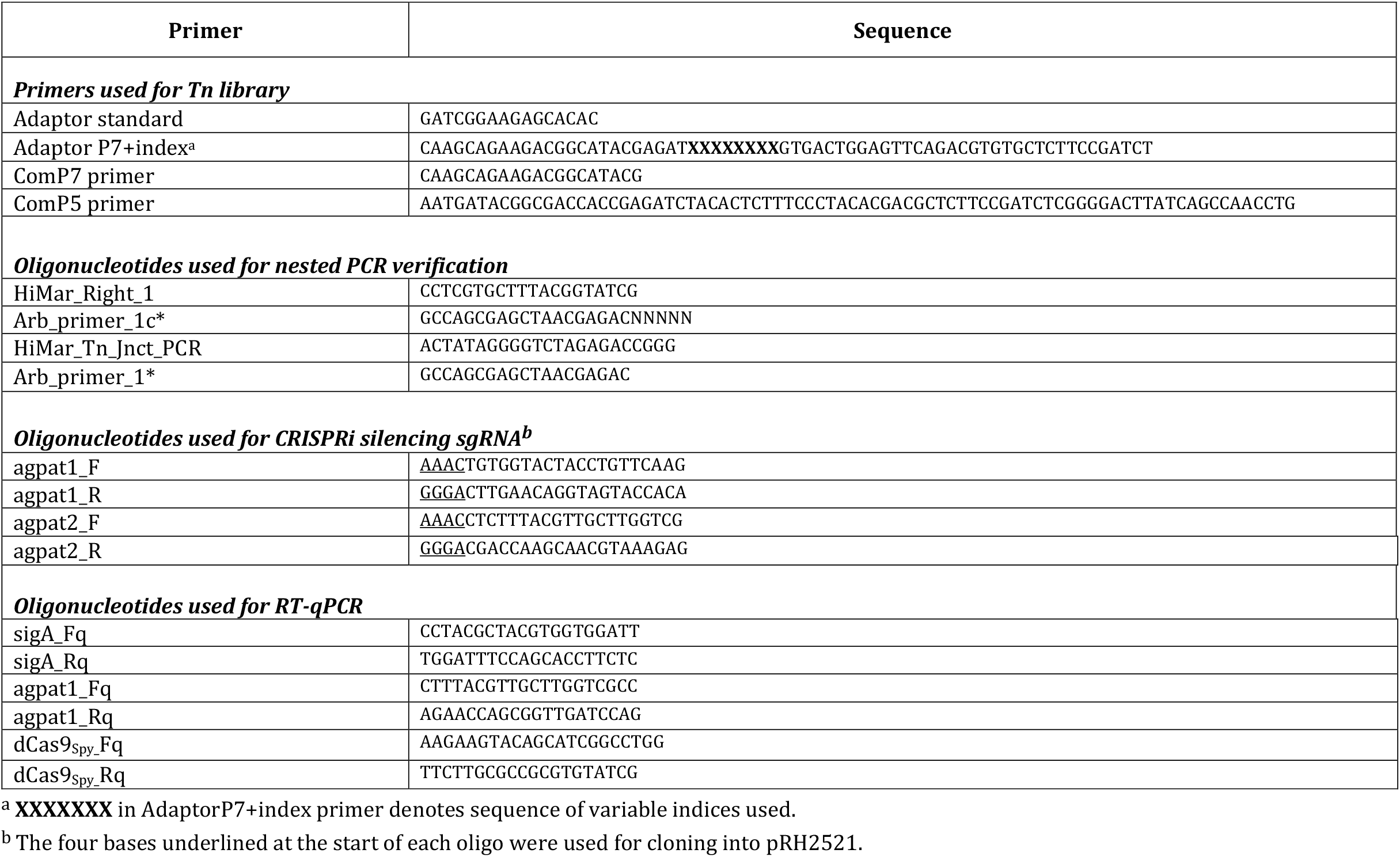
Oligonucleotides used in this study.

**Table 3.** Summary statistics of the Tn libraries created in this study.

|  | *M. bovis*<br>*AF2122/97* | *M. tuberculosis*<br>*H37Rv* |
| --- | --- | --- |
| Unique Mutants | 39,987 (of 73,536) | 29,919 (of 74,604) |
| Saturation | 54% | 40% |
| Essential Genes | 528 | 477 |

### Data Analysis

Indexed libraries were combined, spiked with 20% PhiX, and sequenced on the Illumina HiSeq 3000 platform, using v2 chemistry, generating single-end reads of 150 bp. Raw FASTQ sequence files were pre-processed using the TPP utility of TRANSIT python package (DeJesus *et al*., 2015), including removing TRADIS tags and adapter sequences and mapping using BWA-MEM algorithm [32], to generate insertion files in .wig format. Custom annotations, ‘prot tables’, were created from the *M. bovis* strain AF2122/97 annotation file (NCBI Accession Number LT708304, version LT708304.1) and for the *M. tuberculosis* strain, H37Rv (NCBI Accession Number AL123456, version AL123456.3, assembly build GCA_000195955.2 (ENA)). TRANSIT was run on both *M*.*bovis* and *M. tuberculosis* files using the default normalisation (TTR) and the TRANSIT HMM algorithm (31) to make calls of essentiality for each TA insertion site, and for each gene based on annotated gene boundaries. Data files (fastq) are deposited in SRA (PRJNA754037)

### CRISPRi mediated gene silencing

We utilised dCas9 from *Streptococcus pyogenes* (dCas9_Spy_) for silencing as previously described (32). sgRNA targeting *wag31*_Mtb/Mb_ and *Rv2182c*/*Mb2204c* were designed according to the parameters derived from Larson *et al* 2013 (33). Protospacer adjacent motif (PAM) sites, “NGG,” were chosen and putative sgRNAs 20 bp downstream of the PAM were selected. All sgRNAs designed targeted the coding non-template strand. The probability of complementarity to any other region of the genome and predicted secondary structure of the sgRNA transcript was analyzed using a basic local alignment search tool (BLAST) and M-fold, respectively (34,35). Complementary forward and reverse primers using the sequence (without the PAM) with appropriate ends for ligation to the pRH2521 vector were designed (Table 2). Oligos were annealed and cloned into pRH2521 using BbsI as previous (32,36). One microgram of pRH2502 was electroporated at 25 kV, 25 μF with 1000 Ω resistance into electrocompetent *M. bovis* and *M. tuberculosis* to generate strains expressing dCas9_Spy_ (Mtb__dCas9_/ Mb__dCas9_). These strains were grown and further electroporated with 1 μg of pRH2521 expressing sgRNAs targeting *wag31*_Mtb/Mb_ and *Rv2182c*/*Mb2204c* or pRH2521, the sgRNA -ve plasmid.

### RNA Extraction and RT-qPCR

Cultures were grown to OD_600_ ≅ 0.1–0.2 and the CRISPRi machinery induced with 200 ng/ml of aTc for 1 h. Total RNA was extracted as previously described (37). Briefly, cultures were centrifuged at 3,488 x g at 4°C for 10 min. Pellets were resuspended in 1 ml of TRIzol containing 0.1 mm glass beads and were disrupted by three cycles of 30 sec pulses at 6000 rpm using a Precellys homogenizer. RNA was purified using a Qiagen RNeasy kit combined with on-column DNase digestion according to the manufacturer’s instructions. Quantity and quality were determined using a DeNovix Spectrophotometer (DeNovix Inc, USA) and agarose gel electrophoresis.

To remove traces of contaminating DNA, RNA samples were treated with RNase-free DNase I (Invitrogen) according to the manufacturer’s instructions. cDNA was synthesized from 100 ng of RNA using Superscript III Reverse transcriptase according to manufacturer instructions. qPCRs were performed using PowerUp SYBR Green Master Mix with 1 μl of cDNA and 0.3 μM of either *sigA* primers or gene specific primers (Table 2) in a final volume of 20 μl. Samples were run on a BioRad CFX96 analyser at 50 °C for 2 min, 95 °C for 2 min, followed by 40 cycles of 50 °C for 2 min, 95 °C for 2 min, followed by 40 cycles of 95 °C for 15 sec, 72 °C for 1 min and 85 °C for 5 sec at which point fluorescence was captured. A melt curve analysis was also carried out for each run at 65 °C – 95 °C in increments of 0.5 °C. Gene expression data was analysed using the 2-ΔΔCT method (38). Reverse transcriptase -ve samples were used as a control to ensure removal of gDNA. All results were normalised against the house keeping gene *sigA*. Two or three biological replicates were run, with each measured in duplicate, unless otherwise stated.

## Results

### High-density transposon libraries in *M. bovis* AF2122/97 and *M. tuberculosis* H37Rv were generated

The Mycomar transposon inserts randomly into TA sites in bacterial genomes. There are 73,536 and 74,604 TA sites present in the *M. bovis* (AF2122/97) and *M. tuberculosis* (H37Rv) genomes, respectively. The smaller number of TA sites in *M. bovis* is likely to be reflective of a smaller genome. We successfully generated transposon libraries in *M. bovis* and *M. tuberculosis* containing 39,987 (*M. bovis*) and 29,919 (*M. tuberculosis*) unique mutants, representing 54 % (*M. bovis*) and 40 % (*M. tuberculosis*) saturation. The distribution of transposon insertions in the two species is shown in figure 1.

**Figure 1:**
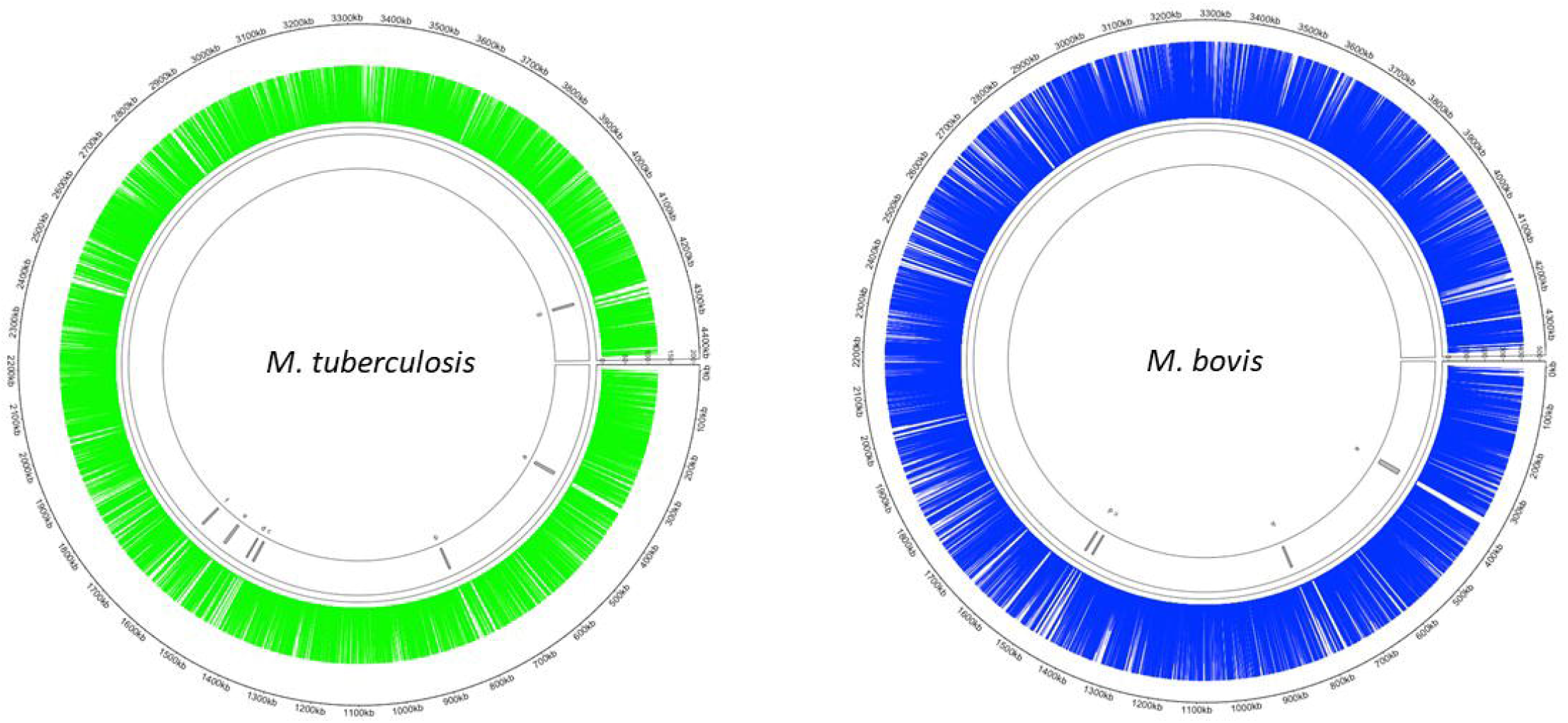
Distribution of Tn insertions in both *M. tuberculosis* and *M. bovis*. Transposon libraries were created in *M. tuberculosis* and *M. bovis* using the *Himar1* system and sequenced on a HiSeq NGS platform (Illumina, UK) as described in the materials and methods. Insertion locations of *Himar1* across the *M. tuberculosis* genome (green) and *M. bovis* genome (blue) were visualised using Circlize (48).

Himar1 transposase has been previously suggested to exhibit local sequence preferences rendering ∼9% of possible TA sites non-permissive to insertion (23) and others have also observed TA insertion cold spots within the *M. tuberculosis* genome. Using the non-permissive sequence pattern, ‘SGNTANCS’ (where S is either G/C), we identified 6657 sites in both *M. bovis* and *M. tuberculosis* genomes (data not shown). Taking a similar approach to Carey *et al*, we found that removing these sites prior to determining gene essentiality as described below did not affect the gene calls (25).

### Comparisons of essentiality between *M. bovis* and *M. tuberculosis*

We examined *in vitro* gene essentiality in *M. bovis* and *M. tuberculosis* using the TRANSIT HMM method (31). This approach classifies genes into four categories; those that are essential for growth and cannot sustain a transposon insertion (ES), those where the transposon insertion results in a growth defect (GD) and those where the transposon insertion results in a growth advantage (GA). Those that show no impact as a result of the transposon insertion are considered non-essential (NE). From this analysis, 527 genes were classified as ES (15.3%), 176 genes were classified as GD (5.1%) and 131 as GA (3.8%) in the *M. bovis* genome. In *M. tuberculosis* 477 genes were classified as ES (13.7%), 179 genes were classified as GD (5.1%) and 1 gene as GA (0.03%). A complete list of calls for the genes that are conserved between both species is given in supplementary table S1. The status of the genes that are *M. bovis* specific are also included in the table.

Early sequencing and functional annotation of the genome of *M. tuberculosis* categorised genes into several different functional classes with an uneven distribution of genes across the classes (4,5). We examined the distribution of the genes classified as ES in *M. tuberculosis* (477) and *M. bovis* (527) across the functional classes to determine if (i) ES genes are over-represented in any particular functional class when compared to the genome as a whole (ii) there are differences between the two species. The results are shown in figure 2 and table 4. Chi squared testing showed that the distribution of ES genes across the functional classes was significantly different to the distribution of all orthologs (p=<0.01). ES genes in both species are over-represented in “information pathways” and “intermediary metabolism and respiration” and under-represented in “conserved hypotheticals” and “PE/PPE” functional classes. Our data are in line with previous reports; Griffin *et al* noted that the distribution of ES genes across the different functional classes were different compared to the genome as a whole (22). DeJesus *et al* also noted that insertions in PE/PPE genes were under-represented likely due to GC rich sequences and an increased proportion of non-permissive TA sites in the PE/PPE genes (23). There were no major differences in distribution of ES genes across the functional classes when *M. tuberculosis* and *M. bovis* were compared with each other except for “insertion sequences and phages” which did not contain any genes classified as ES in the *M. bovis* genome.

**Figure 2:**
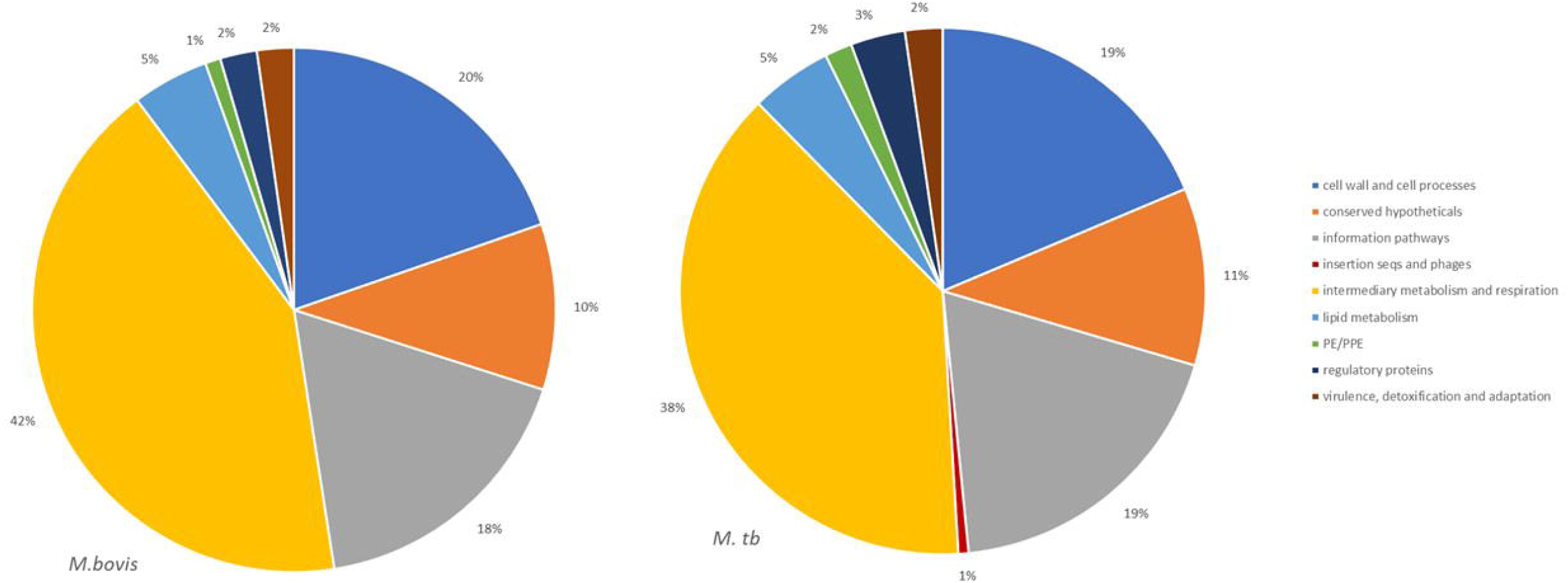
Functional category distribution. Gene essentiality was determined for *M. bovis* and *M. tuberculosis* using custom HMM analyses with TRANSIT software. Functional categories were assigned to orthologous genes and compared for *Himar1* insertion distribution between *M. bovis* (left) and *M. tuberculosis* (right). Transposon insertions were found to be similar across functional categories. Data were analysed using pivot tables in Excel.

**Table 4.**
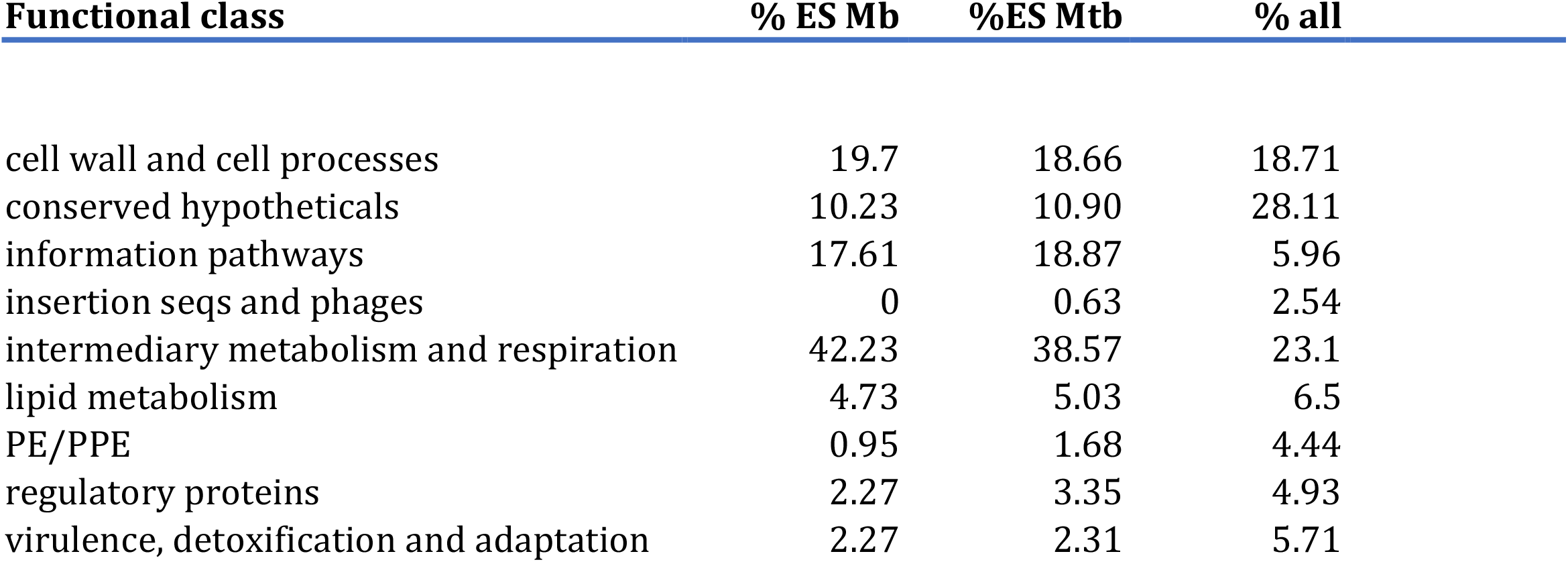
Distribution of genes classified as ES across functional class.

| Functional class | % ES Mb | %ES Mtb | % all |
| --- | --- | --- | --- |
| cell wall and cell processes | 19.7 | 18.66 | 18.71 |
| conserved hypotheticals | 10.23 | 10.90 | 28.11 |
| information pathways | 17.61 | 18.87 | 5.96 |
| insertion seqs and phages | 0 | 0.63 | 2.54 |
| intermediary metabolism and respiration | 42.23 | 38.57 | 23.1 |
| lipid metabolism | 4.73 | 5.03 | 6.5 |
| PE/PPE | 0.95 | 1.68 | 4.44 |
| regulatory proteins | 2.27 | 3.35 | 4.93 |
| virulence, detoxification and adaptation | 2.27 | 2.31 | 5.71 |

Genes categorised as ES in this study were compared between the two species and also compared to previously reported studies (21–23,26,39) (supplementary file, table S1). We found that the *M. bovis* dataset generated in our study shared 370 (70%) of genes classified as ES with *M. tuberculosis in vitro* (this study; figure 3A) and up to 86% overlap with three key published *M. tuberculosis* data sets: DeJesus *et al* 2017 (71%), Griffin *et al* 2011 (86%) and Minato *et al* 2019 (79%) indicating good correlation with previous reports (figure 3D). Similarly, the *M. tuberculosis* dataset generated in our study shared good overlap with other published datasets (figure 3C). When comparing *M. bovis* genes classified as ES with those reported by Butler *et al* 2020 (40) we found that 220 (42%) genes were shared between these data sets (figure 3B). Butler *et al* reported a total of 318 genes to be essential in *M. bovis in vitro* prior to selection in *Dictyostelium discoideum* compared to 527 reported in this study. Both libraries showed similar saturation levels (58% vs. 54% in this study) therefore differences might be due to the conditions under which the libraries were generated (although both studies used Middlebrook 7H11 solid medium supplemented with lysed sheep blood, heat inactivated foetal bovine serum and OADC) or between laboratory variation as might be expected for whole genome techniques such as Tnseq. It should also be noted that the similarities between the studies increases when GD genes are considered, for instance of the 307 genes that appear to be uniquely ES in our study, 212 of these are classified as GD in the study by Butler *et al*., indicating a debilitating impact of the transposon insertion.

**Figure 3:**
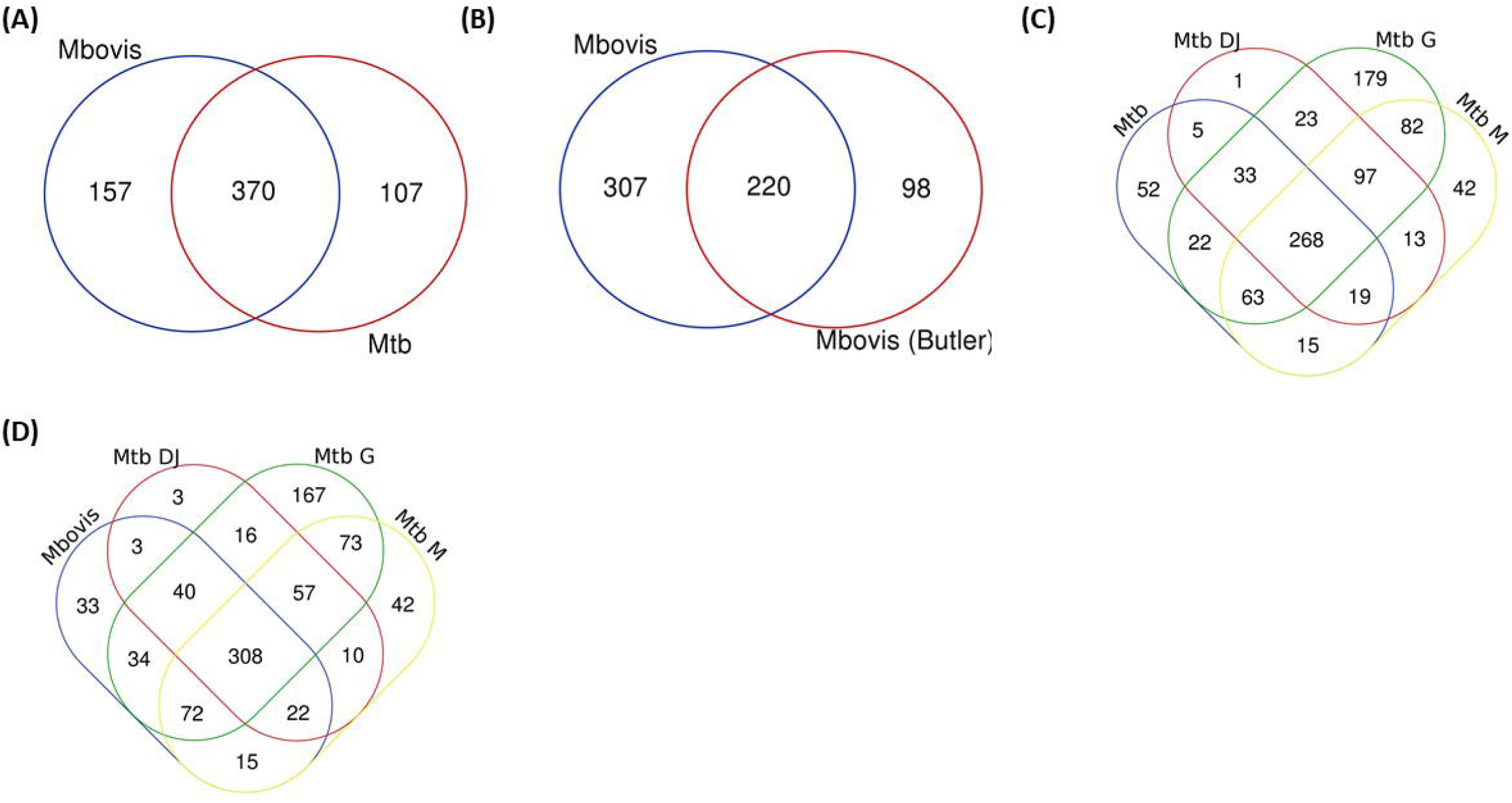
Essential Gene Comparisons. Gene essentiality was determined for *M. bovis* and *M. tuberculosis* using custom HMM analyses with TRANSIT software and compared to previously published datasets. **(A)** *M. bovis* and *M. tb* (both this study), **(B)** *M. bovis* (this study) and *M. bovis* (Butler *et al* 2020) **(C)** *M. tb* (this study) and *M. tb* DJ (DeJesus *et al* 2017), *M. tb* G (Griffin *et al* 2011) and *M. tb* M (Minato *et al* 2019) and **(D)** *M. bovis* (this study) and *M. tb* DJ (DeJesus *et al* 2017), *M. tb* G (Griffin *et al* 2011) and *M. tb* M (Minato *et al* 2019).

### Differences in gene essentiality between *M. bovis* and *M. tuberculosis*

Genes uniquely classified as ES in either species are of interest to determine potential genetic insights for phenotypic differences between these closely related mycobacterial species. In this study 157 genes were uniquely ES in *M. bovis* when compared to the *M. tuberculosis* (figure 3A), however, of these 157, 61 were classified as GD in *M. tuberculosis*. The remaining 96 were classified as NE in *M. tuberculosis* (supplementary file, table S2). The existence of multiple datasets allows for a robust meta-analysis and so we compared across datasets and found that there were 42 genes that were either ES or GD in this study and the study by Butler *et al*., and were classified as NE in *M. tuberculosis* in this study and the study by DeJesus et al. (supplementary file, table S3). Included in this subset of genes is *Rv3543c* (*fadE19*), *Rv3541c* and *Rv3540c* (*lpt2*), genes which are encoded on the same operon (*Rv3545c-Rv3540c* - based on intergenic gaps) regulated by *kstR* and involved in cholesterol catabolism. This study and the Butler *et al*., study indicates that insertional mutagenesis of this operon has a debilitating impact in *M. bovis* but not in *M. tuberculosis*.

Data for the entire *kstR* regulon is given in supplementary file, table S4. Interestingly, the media used in this study and the study by Butler *et al*., contains traces of cholesterol due to the presence of lysed sheep blood, although there is no evidence that cholesterol presented a selective pressure (for *M. tuberculosis*) in this study as there is little overlap of the *M. tuberculosis* dataset with the study by Griffin *et al*., In addition to the *Rv3545c-Rv3540c* operon considered above, several orthologs in the *kstR* regulon were classified as ES in *M. bovis*; *Mb3538* (*Rv3508*), *Mb3568* (*Rv3538*) and *Mb3581* (*Rv3551*), and *Mb3595* (*Rv3565*). Others such as *Mb3541* (*Rv3511*) and *Mb3574c* (*Rv3544c*) were classified as GD. Interestingly insertions in the genes belonging to the *mce4* operon and required for growth on cholesterol mostly confer a GA for *M. bovis*. These observations might reflect a difference in the requirement for cholesterol catabolism *in vitro* in a complex carbon mixture compared to *M. tuberculosis*.

One of the key metabolic differences between *M. bovis* and *M. tuberculosis* is the inability of *M. bovis* to utilise carbohydrates. Genes in the glycolytic pathway (supplementary file, table S5) such as, enolase (*eno*), pyruvate kinase (*pykA)* and pyruvate carboxylase (*pca*) might be expected to be NE in *M. bovis* as *pykA* is non-functional in *M. bovis* (41). The datasets show that *eno* is ES in *M. bovis* as well as *M. tuberculosis* perhaps indicating that its essentiality is linked to a role other than glycolysis. Similarly, the suggestion that a transposon insertion in *pykA* confers a GA (this study only) is counter-intuitive and might suggest a non-glycolytic role for this enzyme. Only our dataset suggests that a transposon insertion in *icl1*, an enzyme required for growth on fatty acids, confers a growth advantage in *M. bovis*.

The two-component system PhoPR has been shown to regulate *de novo* PDIM synthesis and also co-ordinate the acid-stress response (16,42). It is of particular interest because a non-synonymous SNP in the sensor histidine kinase *phoR* in *M. bovis* renders signalling through the system defective, however, the existence of compensatory mechanisms that restore PDIM synthesis obscures the role of the regulon in *M. bovis*. Of the genes in the PhoPR regulon (supplementary file, table S6) only *Rv3778c* seems to be consistently required across species and studies. Genes in the redox sensing WhiB family are included in the operon (*whiB1, whiB3* and *whiB6*) but only *whiB1* is ES in *M. bovis* in our study.

Finally, as the electron transport chain and ATP synthesis is a relatively new therapeutic pathway we chose to examine ES more closely in these pathways (supplementary table, S7). These pathways are targets of recently introduced drugs such as bedaquiline (ATP synthase) and those in development e.g. Q203 which targets the terminal cytochrome bc_1_-aa_3_ oxidase (43). Unsurprisingly, the genes encoding the ATP synthase are largely ES in both species in all studies (*Rv1304-Rv1311*) with the exception of *Rv1304* (*atpB*). The genes that encode a sub-unit of the terminal cytochrome bc1-aa3 oxidase complex (*qcrCAB*) the target of Q203 are classified as either ES or GD. One interesting observation is that both our study and the study by Butler *et al*., is that a GD occurs as a result of an insertion in *nuoG* but this is not observed in any of the *M. tuberculosis* studies. *nuoG* forms part of the multi-subunit NADH reductase-I complex in the respiratory chain and transfers electrons to the menaquinone pool while simultaneously contributing to the proton gradient through its proton pumping function

### Establishment of CRISPRi in *Mycobacterium bovis* using *wag31*

Wag31 is required for peptidoglycan synthesis and previously published datasets have classified *wag31* in *M. tuberculosis* as ES *in vitro* (21–23). This study classified *wag31* in *M. bovis* as ES but NE in *M. tuberculosis*. The study by Butler *et al*., assigned *wag31* as NE in *M. bovis*. In order probe this discrepancy and to establish CRISPRi silencing in *M. bovis* this gene was chosen for silencing. Early CRISPRi studies in *M. tuberculosis* performed by Singh *et al*. successfully utilised two plasmids encoding sgRNAs guides targeting +26 bp to +45 bp and +144 bp to +163 bp downstream of the annotated start codon of *wag31*_Mtb_ (table 1 and figure 4A). We utilised these plasmids to investigate the impact of silencing *wag31*_Mb_. *M. bovis* AF2122/97 was transformed with pRH2502 to create a strain expressing *dcas9*_*Spy*_ (Mb__dCas9_). Mb__dCas9_ was then transformed with plasmids expressing the sgRNA guides. Strains were cultured to exponential phase and serial dilutions were spotted onto agar containing 200 ng/ml aTc. Controls (without aTc, without sgRNA) were also included. The results, presented in figure 4B, show that silencing *wag31*_*Mb*_ in *M. bovis* results in a severe growth defect, visible at 10^−1^ dilution with complete cessation of growth at 10^−2^ dilution. This growth defect is identical to that seen in *M. tuberculosis* and supports the ES classification.

**Figure 4.**
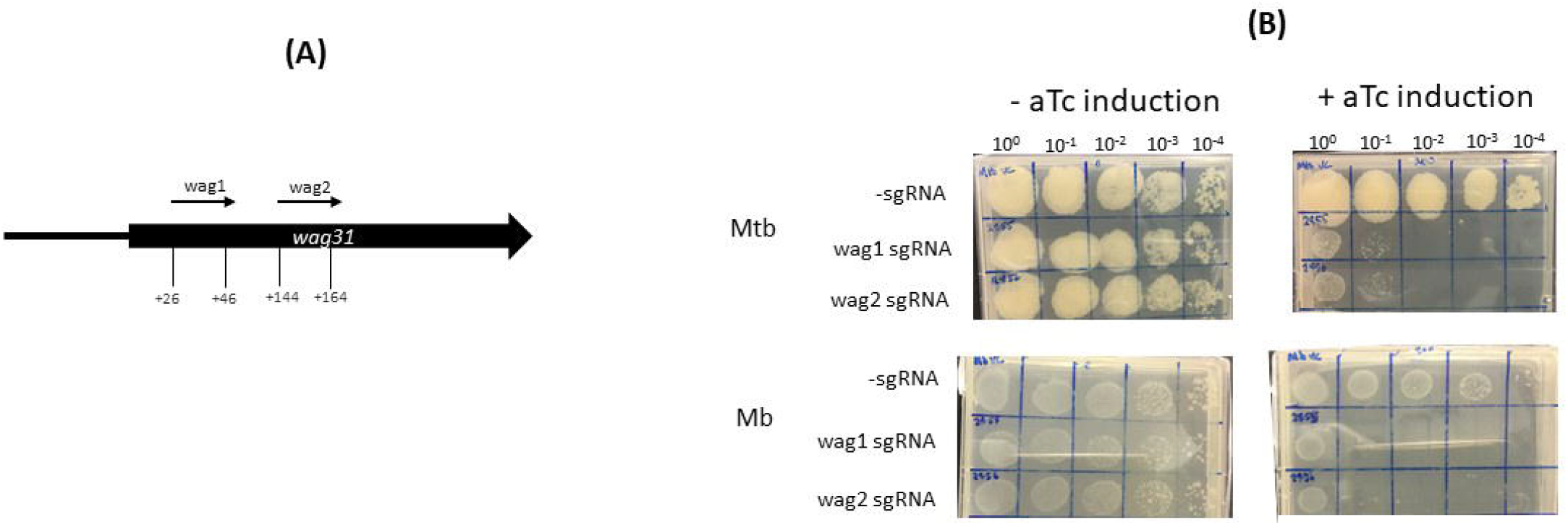
Using CRISPRi/dCas9 to inhibit *wag31* expression in *M. bovis* and *M. tuberculosis*. **(A)** Schematic showing the target regions of two sgRNAs designed to target and inhibit *wag31* expression. The numbers show the nucleotide position of the sgRNA relative to the annotated start site **(B)** CRISPRi strains were cultured in 10 ml of supplemented 7H9 medium to exponential phase and diluted to 2 × 10^7^ CFU/ml. A 10-fold serial dilution to 10^−4^ was performed and 20 μl of each dilution was spotted onto 7H11 agar without aTc and with 200 ng / ml aTc to induce CRISPRi/ dCas9 and the sgRNA in those strains that carried the guide. Two biological replicates were carried out.

### Silencing *Rv2182c* and its ortholog *Mb2204c* shows a species-specific growth impact

*Rv2182c*/*Mb2204c* is annotated as a 1-acylglycerol-3-phosphate O-acyltransferase (agpat) and involved in glycerophospholipid metabolism. It is thought to synthesize diacylglycerol-3P through the addition of acyl chains to monoacylglycerol-3P. It is classified as ES in *M. tuberculosis* in this study and by others (22,23,44). It is classified as ES in *M. bovis* in this study but NE in the study by Butler *et al*.,. Strains of *M. tuberculosis* and *M. bovis* were constructed expressing sgRNAs targeting +2 bp to +21 bp and +40 bp to +59 bp downstream of the annotated start codon of *Rv2182c*/*Mb2204c* (table 2 and figure 5A). The impact of inducing the system on expression of *Rv2182c/Mb2204c* was measured using RT-qPCR. The results, which are shown in figure 5B show that dCas9_Spy_ is similarly induced in both *M. tuberculosis* and *M. bovis* with 150 to 350-fold induction of expression in the presence of aTc. Additionally, the results show that, in the presence of the sgRNA, there is a clear reduction in expression of *Rv2182c/Mb2204c* in both species. These data demonstrate effective gene silencing of *Rv2182c/Mb2204c* in both *M. tuberculosis* and *M. bovis*, respectively.

**Figure 5.**
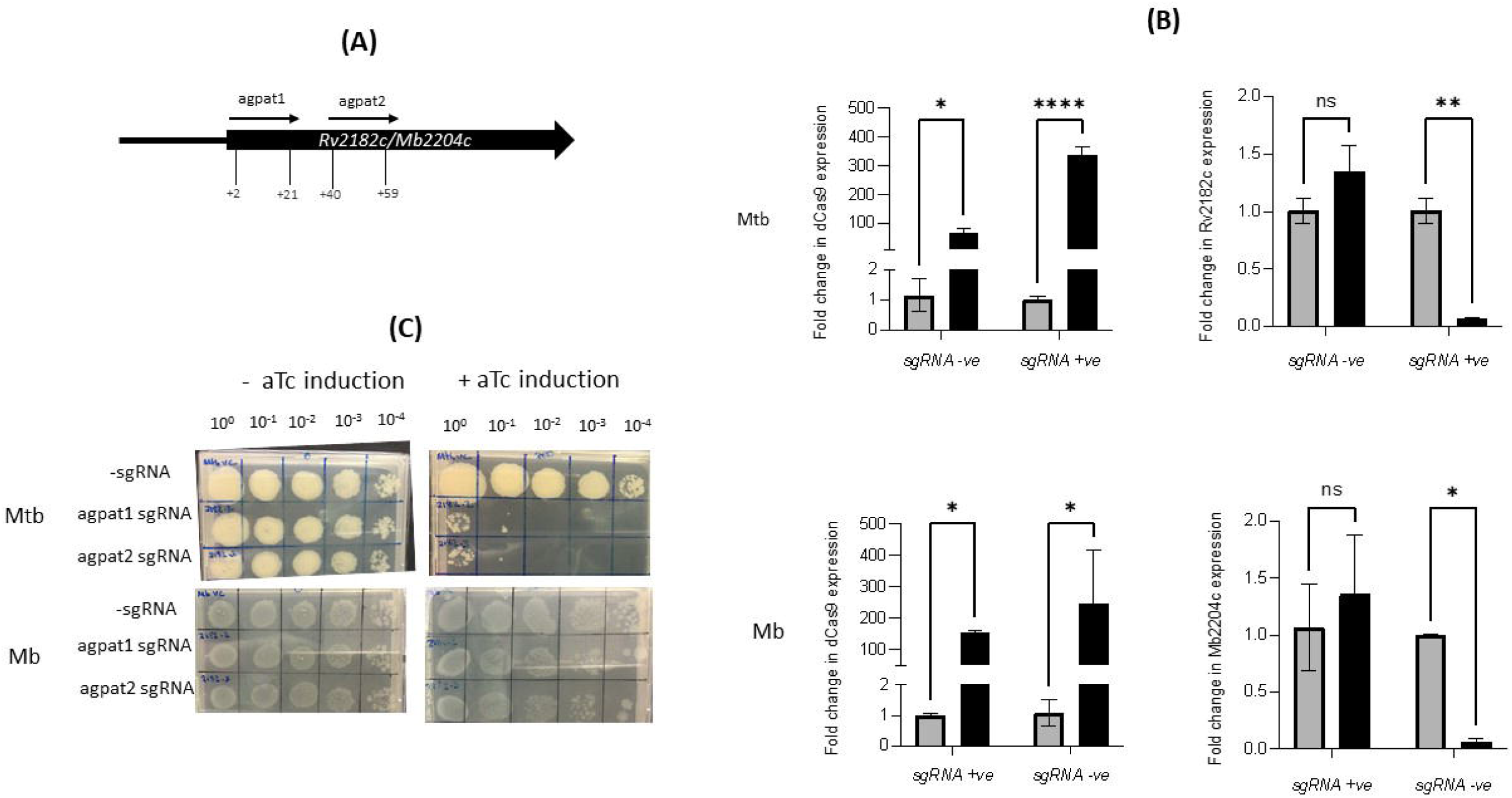
Using CRISPRi/dCas9 to inhibit *Rv2182c/Mb2204c* expression in *M. bovis* and *M. tuberculosis*. **(A)** Schematic showing the target regions of two sgRNAs designed to target and inhibit *Rv2182c/Mb2204c* expression. The numbers show the nucleotide position of the sgRNA relative to the annotated start site **(B)** *dCas9* expression and *Rv2182c/Mb2204c* expression were measured by RT-qPCR as described in the methods section. Gene expression was analysed using the 2^-ΔΔCT^ method, normalised against *sigA*. Results represent two biological repeats with two technical repeats each. *P* = * <0.05, ** <0.01, *** <0.001, **** <0.0001 or not significant (ns), analysed using a 2-way ANOVA test **(C)** CRISPRi strains were cultured in 10 ml of supplemented 7H9 medium to exponential phase and diluted to 2 × 10^7^ CFU/ ml. A 10-fold serial dilution to 10^−4^ was performed and 20 μl of each dilution was spotted onto 7H11 agar without aTc and with 200 ng/ml aTc to induce CRISPRi/ dCas9 and the sgRNA in those strains that carried the guide. Two biological replicates were carried out.

To determine the impact of induction of the guides, strains were cultured to exponential phase and serial dilutions were spotted onto agar containing 200 ng/ml aTc. The results (figure 5C) show that silencing *Rv2182c* in *M. tuberculosis* results in a severe growth defect, with almost complete cessation of growth at 10^−1^ dilution. However, the consequence of silencing *Mb2204c* on the growth of *M. bovis* is far less impactful with a small reduction visible at the lowest dilution 10^−4^. This demonstrates that, unlike *wag31*, silencing of *Rv2182c* and its ortholog *Mb2204c* in *M. tuberculosis* and *M. bovis* respectively, has a differential impact on growth, with *M. tuberculosis* being more vulnerable and showing a greater growth defect. These results do not support the classification of *Mb2204c* as an ES gene in *M. bovis* but they clearly highlight that there are different phenotypic consequences as a result of silencing the ortholog in both species.

## Discussion

The aim of this work was to directly compare gene essentiality in the human and animal adapted members of the MTBC. In order to do this we generated transposon libraries in *M. bovis* AF2122/97and *M. tuberculosis* H37Rv using a rich medium that supported the growth of both species. We assessed gene essentiality using the TRANSIT HMM method to define 527 and 477 genes as ES for *M. bovis* and *M. tuberculosis*, respectively. Datasets from each species were compared with each other and with previously published datasets. Genes classified as ES were congruent between the species and also with existing studies of gene essentiality in *M. tuberculosis* (21–23,39). Comparing this study with a previously published *M. bovis* dataset revealed a 42% overlap which increased when genes predicted to show a GD as a result of the transposon insertion were taken into account (26). There were some indications of differences between the species, and a meta-analysis of the data indicated that 42 genes were differentially essential between the species. A recent study using whole genome CRISPRi screens showed that a similar number (80 genes) were differentially essential in two different strains of *M. tuberculosis* (H37Rv vs HN878) (45). Genes that appear to show differential essentiality between the two species include those involved in cholesterol catabolism.

Whole-genome TnSeq provides a high-throughput assessment of fitness costs and has allowed the classification of genes based on essentiality but does not provide information on target vulnerability. More recent studies highlight the limitations of the (near) binary classification of genes into and ES/NE and utilise CRISPRi to assess vulnerability (45,46). Additionally, datasets are prone to false calls of ES due to non-saturating mutagenesis. In this study CRISPRi was utilised to show that there are different impacts on bacterial growth as a result of silencing *Rv2182c*/*Mb2204c* in their respective species, despite achieving similar levels of gene silencing. Significant growth inhibition was seen as a result of silencing in *Rv2182c* in *M. tuberculosis* while only marginal impacts on growth were observed on silencing the ortholog *Mb2204c* in *M. bovis. Rv2182c*/*Mb2204c* is annotated as a 1-acylglycerol-3-phosphate O-acyltransferase and involved in glycerophospholipid metabolism. It is thought to synthesize diacylglycerol-3P through the addition of acyl chains to monoacylglycerol-3P. This pathway may be involved in detoxification and further work is required to fully understand the differential impact of silencing this gene in the two species. Given that *Rv2182c* was a predicted target in a recent compound screen (47), differential essentiality estimates in *M. bovis* and *M. tuberculosis* are important to predict if zoonotic TB caused by *M. bovis* can also be suitably treated with drugs designed to be effective against *M. tuberculosis*.

We have provided a comparative analysis of the genetic requirements for growth of two key MTBC members: *M. bovis* and *M. tuberculosis*. Genes which are uniquely ES for either *M. bovis* or *M. tuberculosis* have the potential to provide insights into niche specific aspects e.g., host tropism, survival in the environment, phenotype, and anti-tubercular drugs. Host tropism is of particular interest when considering the zoonotic nature of *M. bovis* and the involvement of wildlife hosts as reservoirs of infection for bovine TB. Use of *M. bovis* libraries in the context of the host i.e., through experimental infection of bovine TB will enable the study of the genetic requirements for survival *in vivo*. Further investigations exploring the role and function of ES genes between *M. bovis* and *M. tuberculosis* is necessary to better understand the physiological differences in these key MTBC species.

## Supporting information

Supplemental Methods and Results S2

## Author Contributions

AJG, SW, IP and SLK designed the study. AJG, IP, VF, carried out the experimental work. Data analysis was done by IN, JS and DX. SLK, DW, BWW and BVR did funding acquisition. AJG and SLK wrote the first draft of the manuscript. All authors contributed to the manuscript revision, read, and approved the submitted version.

