## Supplemental Methods and Results S2 for "Probing differences in gene essentiality between the human and animal adapted lineages of the *Mycobacterium tuberculosis* complex using TnSeq"

**Supplementary Methods and Results**

1. **Validation of Random Transposon Insertion**

Transduced *M. bovis* Transposon libraries were recovered on selective modified 7H11 medium containing 25 µm/ml Kanamycin for up to 6 weeks. Individual mutants were selected and used to inoculate 10 ml of in 7H9 medium containing 75 mM sodium pyruvate, 0.05% Tween^®^80 and 10% ADC and cultures were incubated at 37°C until reaching 0D_600_ ≅ 0.8. Genomic DNA was extracted from 1 ml of culture by bead beating followed by Phenol-Chloroform extraction and Ethanol precipitation as described in main materials and methods.

A nested PCR approached was used followed by Sanger sequencing to determine the Transposon insertion site **(Figure 1.1A)**. Briefly the process order is as follows: PCR round #1, DNA clean up, PCR round #2; assess by gel electrophoresis, PCR clean up, Sequence.

*Primers:*

HiMar_Right_1	CCTCGTGCTTTACGGTATCG

Arb_primer_1c*	GCCAGCGAGCTAACGAGACNNNNN (random primer)

HiMar_Tn_Jnct_PCR	ACTATAGGGGTCTAGAGACCGGG

Arb_primer_1*	GCCAGCGAGCTAACGAGAC

*PCR conditions:*

Round #1 Amplification

1. 95 °C for 5 min
2. 95 °C for 1 min
3. 38 °C for 1 min
4. 72 °C for 2 min – go to step 2 x 30 cycles
5. 72 °C for 2 min

Round #1 Amplification

1. 98 °C for 30 sec
2. 98 °C for 15 sec
3. 55 °C for 30 sec
4. 72 °C for 30 sec – go to step 2 x 30 cycles
5. 72 °C for 2 min

Analysis of Round #2 products by sequencing provides the location of the Tn within the genome. From this position specific PCR probes ~1 kb into the genome were then designed to verify Transposon location and orientation. PCR products were verified by Sanger sequencing in both directions. Approximately 5 mutants per Tn library were verified in this way. Using a combination of Nested PCR sequencing and specific PCR sequencing the locations of Transposon insertions were mapped to *M. bovis* AF2122/97 genome using Artemis **(Figure 1.1B)**

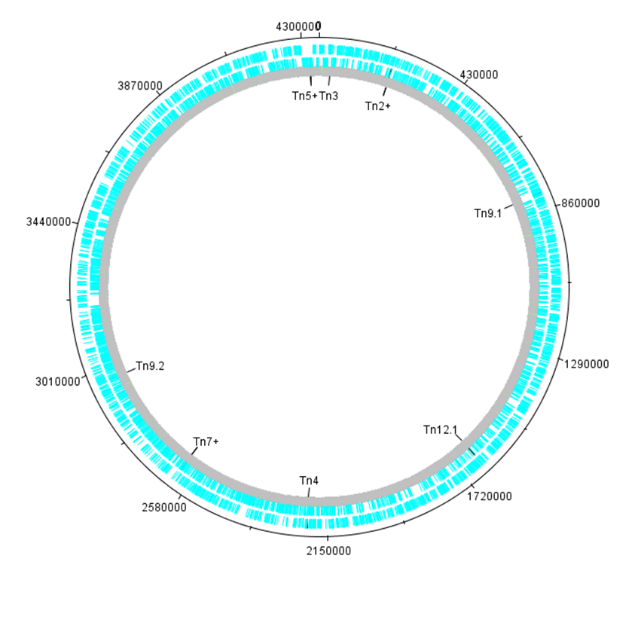

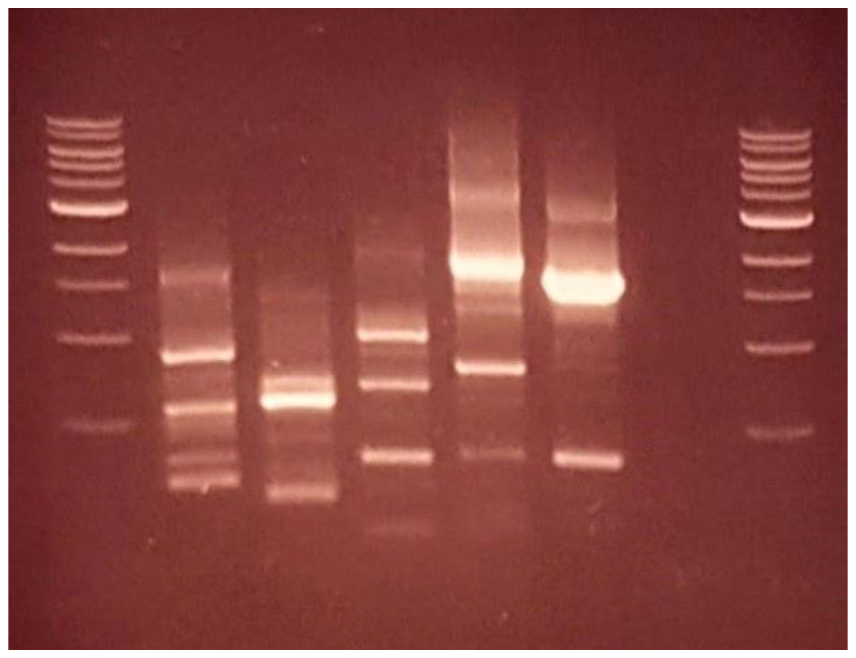

**B**

**A**

1 kb	Tn1	Tn2	Tn3	Tn4	Tn5	Neg	1kb

**Figure 1.1 *M. bovis* Transposon Library Validation**
